# Rapid emergence of hyperparasitic elements may stop *P-element* invasions in the absence of a piRNA-based host defence

**DOI:** 10.1101/2025.03.10.642342

**Authors:** Matthew Beaumont, Divya Selvaraju, Riccardo Pianezza, Robert Kofler

**Author notes:** to whom correspondence should be sent. authors contributed equally (Matthew Beaumont, Divya Selvaraju).

## Abstract

Transposable element (TE) invasions pose risks to both the TE and the host. All copies of a TE may be lost via genetic drift, or host populations may suffer fitness declines, potentially leading to extinction. By monitoring invasions of the *P-element* in experimental *D. melanogaster* populations for over 100 generations, we uncovered a novel risk for invading TEs. In two replicate populations, the *P-element* rapidly multiplied until a piRNA-based host defence emerged, leading to the plateauing of TE copy numbers. However, in one population (R2), *P-element* copy numbers stabilised at a significantly lower level, despite the absence of a piRNA-based host defence. We find that this stabilisation was likely driven by the propagation of non-autonomous insertions, characterised by internal-deletions, which out-competed the autonomous full-length insertions. Such a rapid proliferation of non-autonomous insertions could account for the high prevalence of *P-element* insertions with internal-deletions observed in natural *D. melanogaster* populations. Our work reveals that TEs may stochastically sabotage their own spread in populations due to the emergence of hyperparasites, rendering the establishment of a host defence unnecessary. The proliferation of hyperparasitic elements may also lead into an evolutionary dead end, where affected populations are resistant to re-invasion (e.g. following recurrent horizontal transfer), yet are unable to infect other species due to a lack of autonomous insertions.

## Introduction

Eukaryotic organisms have long faced the threat of transposable element (TE) invasions. These stretches of DNA integrate into host genomes and selfishly replicate, irrespective of fitness effects [15, 49]. TEs have proven extraordinarily effective in self-transmission, having been able to invade almost all observed eukaryotic genomes [5, 75]. They show varying success rates in colonising different species, where the total TE content in host genomes ranges from just 3% in yeast, to 78% in Antarctic krill [5, 66]. Although some TE insertions have been posited to be beneficial to a host [2, 20], it is generally assumed that most insertions are either neutral or deleterious. Left unchecked, the ever-propagating TE poses a threat to genome stability and potentially even population survival as a whole [48, 31, 32]. In response, hosts have developed intricate defence mechanisms by which they can limit replication, typically utilising small RNAs [61]. In *Drosophila*, host defence is based around piRNAs, small RNAs ranging in size from 23 to 29 nt that silence TEs at both the transcriptional and post-transcriptional levels [22, 9, 68, 38]. These piRNAs are derived from distinct genomic loci, termed piRNA clusters, comprising around 3.5% of the total genome in *D. melanogaster* [78]. A fundamental component of piRNA biogenesis is the ping-pong cycle, which amplifies the abundance of piRNAs through a positive feedback loop involving two cytoplasmic proteins, Aub and AGO3 [9, 22]. Cleavage of TE transcripts by Aub yields novel piRNAs, which may then be loaded into AGO3 to guide the cleavage of further transcripts into piRNAs, that are then again loaded into Aub. Slicing of TE transcripts also brings about ‘phasing’, in which the resulting piRNA precursors are processed by the endonuclease, Zuc [23, 14]. Whilst the ping-pong cycle amplifies piRNA abundance, phasing is thought enhance piRNA diversity [23, 46, 14]. It is unclear what initially triggers the emergence of a piRNA-based host defence [41, 62, 19]. The current prevailing model, the trap-model, holds that an invading TE is stopped when it transposes into a piRNA cluster, which then triggers the production of piRNAs that suppress the TE [4, 43, 79, 50].

Another mechanism that may affect the proliferation of a TE, are non-autonomous insertions. Non-autonomous elements were first described by Barbara McClintock as the now famous pair of loci, Ds and Ac, controlling chromosomal breakage in maize [45]. Later, it was discovered that Ac is a full-length autonomous TE and Ds a non-autonomous element, whose activity depends on Ac [18]. These non-autonomous insertions are unable to produce the proteins necessary for mobilisation, but can utilise the proteins generated by autonomous insertions [25]. Therefore, they have been described as hyperparasites, able to take advantage of the molecular machinery of the autonomous full-length insertions. Non-autonomous insertions have been observed for many TE families [25]. It is possible that some non-autonomous elements benefit from a mobilisation advantage over autonomous insertions. For *Mariner* in *D. melanogaster*, the non-autonomous element (*Peach*) seems to proliferate more efficiently than the autonomous (*Mos1*) [59]. A notable example of these dynamics, and focus of this study, are the hyperparasites of the *P-element* in *Drosophila* species. The *P-element* is a 2907 bp DNA transposon with four ORFs [6] that was famously discovered as the causative agent of hybrid dysgenesis, where the offspring of reciprocal crosses among the same two strains may show varying ovarian phenotypes (i.e. atrophied vs regular) [29, 6]. The *P-element* is active in the germline but not in the soma [36], theorised to be a strategy to minimise damage to the host [11]. This tissue specificity is regulated by alternative splicing of the third intron (IVS3), which is spliced out in the germline but retained in the soma [36]. Interestingly, the piRNA-based host defence acts by repressing IVS3 splicing in the germline [71]. Internal deletions (IDs) of the *P-element* have been frequently observed in prior work [7, 17, 65, 34, 33]. Many of these *P-element* insertions with IDs are non-autonomous and may be mobilised by the transposase produced from full-length insertions. Some may be preferentially mobilised relative to their full-length copies [26, 34, 65]. Interestingly, certain non-autonomous *P-element* insertions, like the *KP*-element or *D50*, may even act as repressors of *P-element* activity [7, 56]. Non-autonomous *P-element* insertions may yield defective transposase proteins that are able to occupy transposase binding sites, thereby blocking access of functional transposases preventing mobilisation [40]. Importantly, the regulation of TEs by non-autonomous elements is not a feature of the host defence but rather a limitation of the mechanism by which TEs replicate. Irrespective of how TE activity is controlled, inactive TE families will gradually accumulate mutations that will render all TE copies non-functional. Such inactive TEs will therefore eventually face extinction [8]. To escape this gradual erosion by mutations, TEs occasionally undergo horizontal transfer (HT) into a novel unprotected species, where they are able to replicate until they are once again silenced by the host. Such TE invasions triggered by HT may be far more common than previously thought [62, 52, 51]. However, the invasion of novel species may pose some risk for the newly arrived TE as well as to the host. First, even after successful HT into a novel species, a TE may fail to become established in a host population [37]. All copies of the TE may be lost due to drift [37] or negative selection. Second, TE invasions could dramatically reduce the fitness of the host, such that the survival of the host population is threatened [31, 32]. For example, we previously found that the establishment of the piRNA-based host defence may fail stochastically in populations invaded by the *P-element*, with dramatic effects on host fitness [65]. A startling decline in host fitness that eventually led to the extinction of the experimental population has also been seen in other works [72]. Extinction of host populations will, of course, also all remove all active TEs copies. Assuming that each TE only has but a few opportunities to spread by HT to a novel species before gradual erosion by mutations deactivates all copies, it is crucial for the TE to efficiently utilise these limited opportunities, or otherwise face extinction.

By monitoring *P-element* invasions in three experimental *D. melanogaster* populations for 100 generations, we discovered a novel threat to the long-term persistence of TEs. The *P-element* spread rapidly in two (R1, R3) out of the three replicates, where the emergence of a piRNA-based host defence led to stable copy numbers between generations 30-40. However, in one replicate (R2) copy numbers stabilised at around the same time at a significantly lower level, despite the absence of a piRNA-based host defence. *P-element* insertions with IDs, likely non-autonomous elements, proliferated in this replicate to such an extent that few autonomous copies remained, likely resulting in the stabilisation of *P-element* copy numbers. Our work reveals a novel risk of TE invasions, i.e. that hyperparasites may rapidly emerge during an invasion and spread such that few autonomous copies remain. Such a proliferation of hyperparasitic elements may lead into an evolutionary dead end, endangering the long-term persistence of TEs. With no host defence emerging in R2 during the experiment, our work also raises questions as to what triggers the initiation of a piRNA-based host defence.

## Results

### *P-element* invasions in experimental *D. melanogaster* populations

To study the dynamics of TE invasions, we introduced the *P-element*, via micro-injection, into a *D. melanogaster* strain (DM68) without any *P-element* insertions (supplementary fig. S1). We then established three replicate populations (R1, R2, R3), by mixing transformed flies with näive DM68 flies. Populations were maintained at a size of *N* = 250 and a temperature of 25°C, with non-overlapping generations. We monitored the following *P-element* invasion in each population for more than 100 generations. At regular time intervals, we used short-read sequencing on pooled genomic DNA from each replicate, as well as the transcriptome (Fig 1A). We used the tool DeviaTE [73] to estimate the *P-element* copy number in each sample, which normalises the coverage of TEs to the coverage of single-copy genes (Fig 1C). For example, if the *P-element* has a coverage of 50x and the single-copy genes an average coverage of 5x, then we infer 10 *P-element* copies per haploid genome.

**Figure 1:**
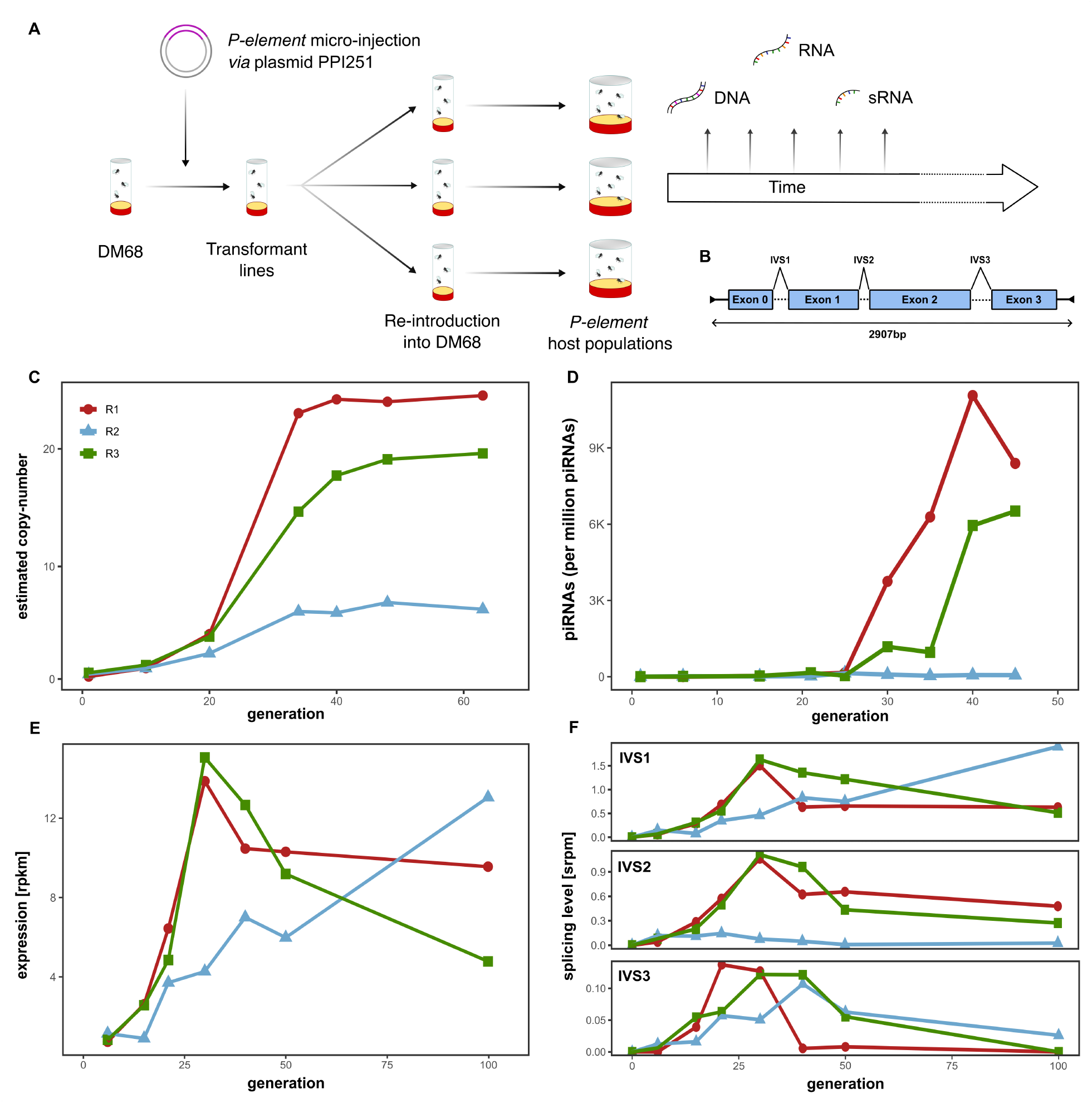
*P-element* invasions in three experimental *D. melanogaster* populations (R1, R2, R3). **A** Overview of our experimental design. **B** Basic schematic of the *P-element*, including exons, intervening sequences (IVS) and TIRs (black arrows). **C** *P-element* copy numbers in each replicate across time. **D** piRNAs mapping to the *P-element* across time. **E** Expression of the *P-element* across time. **F** Abundance of spliced reads for each *P-element* intron (IVS1-3) in spliced reads per million (srpm).

We observed that *P-element* copy numbers increased in all replicates until generations 35-40, where copy numbers reached a stable plateau (Fig. 1C). In R1 and R3, this plateau was around 20-26 copies per haploid genome, whereas in R2, copy numbers plateaued at a much lower level of ∼7 copies. We then asked what could be responsible for the lower level observed in R2. We first tested whether the piRNA-based host defence was established faster in R2 than in the other replicates. A rapid emergence of a host defence could limit the accumulation of *P-element* copies. To assess this, we investigated the abundance of small RNAs mapped to the *P-element* in the experimental populations. Contrary to our expectations, we found that a large number *P-element* piRNAs emerged around generation 25-40 in R1 and R3 but not in R2 (Fig. 1D). In addition, the number of siRNAs was very low in R2 (supplementary Fig. S2). This raises the important question as to how the invasion was so effectively cut short without a piRNA-based host defence. Next, we wondered whether the *P-element* was silenced via another mechanism, independent of piRNAs. To test this, we sequenced the bulk mRNA of each pooled sample. *P-element* expression increased in all replicates until generation 30 (Fig. 1E). In R1 and R3, expression then slowly declined. Contrastingly, *P-element* expression in R2 increased continuously over time, reaching it’s highest value at generation 100 (i.e. the latest available time point; Fig. 1E). Therefore, our data suggests that the *P-element* transcription was not repressed in R2.

Splicing of the *P-element* introns is essential for it’s biology [1, 67]. IVS3 particularly so, as the tissue specificity of the *P-element* is regulated by alternative splicing of this intron (IVS3 is spliced out in the germline but retained in the soma [36]). Additionally, the piRNA-based host defence silences the *P-element* by repressing splicing of IVS3 [71]. We estimated the splicing level of all three introns of the *P-element* during our experiment (Fig. 1F). In R1 and R3, the splicing of all three introns decreased around generation 20-30, where the level of splicing of IVS3 was most dramatically reduced. By contrast, we still found high levels of splicing of IVS3 at generation 100 in R2 (Fig. 1F). Here, splicing of IVS2 also remained at a constantly low level throughout, in contrast to the levels of IVS1 and IVS3 splicing, that either increased or fluctuated during the experiment (Fig. 1F).

Together, these results show that in two replicates (R1 and R3), *P-element* copy numbers increase to approximately 20 copies per haploid genome and then stabilise as piRNAs emerge. This stabilisation in copy numbers coincides with reduced *P-element* expression and splicing of IVS3. In one replicate (R2), copy numbers plateau at a significantly lower level, despite lacking a piRNA-based host defence. *P-element* expression in R2 also continues to increase and IVS3 splicing is still observed after 100 generations.

### Inactive ping-pong cycle in R2

We next investigated in more detail as to why so few *P-element* piRNAs were generated in R2. In R1 and R3, small RNAs mapping to the *P-element* are primarily between 23-29 nt long, and largely have a ‘U’-bias at the first base, as expected of piRNAs (supplementary fig. S3[9]). However, in R2, small RNAs mapping to the *P-element* predominantly have a size of 21 nt, alongside a less pronounced U-bias than both R1 and R3, more indicative of siRNAs than piRNAs (supplementary fig. S3; [14]). At later generations, piRNAs are broadly distributed over the *P-element* in R1 and R3 and almost entirely absent in R2 (Fig. 2A, 1D; supplementary fig. S4). piRNA abundance is thought to be amplified by the ping-pong cycle, a positive feedback loop, wherein the cleavage of sense and antisense transcripts of TEs results in novel piRNAs [9, 39]. Therefore, we questioned whether the ping-pong cycle was inactive for the *P-element* in R2. An active ping-pong cycle leads to a distinct signature; piRNAs produced from opposing strands will frequently overlap by 10 nt at the 5’ ends, termed the ping-pong signature [9, 22]. We found a ping-pong signature emerging for the *P-element* between generations 25-30 in R1 and R3, but we could not observe a ping-pong signature at any generation in R2 (Fig 2B; supplementary fig. S5). However, the ping-pong cycle is functional in R2, as we found a clear signature for another TE (Blood) (supplementary fig S6). Downstream of the ping-pong cycle, an additional process termed ‘phasing’ may be active, in which cleaved piRNA precursors are further processed into piRNAs by the endonuclease Zucchini [23, 46, 14]. Phasing also leads to a characteristic pattern in the distribution of the distance between the 5’-end and 3’-start of neighbouring piRNAs, where a distance of 1 nt is overrepresented [23]. We observe this signature for *P-element* mapping piRNAs at later generations in R1 and R3, but have too few piRNAs to compute it in R2 (supplementary fig. S7). To summarise, the absence of the ping-pong cycle for the *P-element* likely accounts for the low abundance of piRNAs in R2.

**Figure 2:**
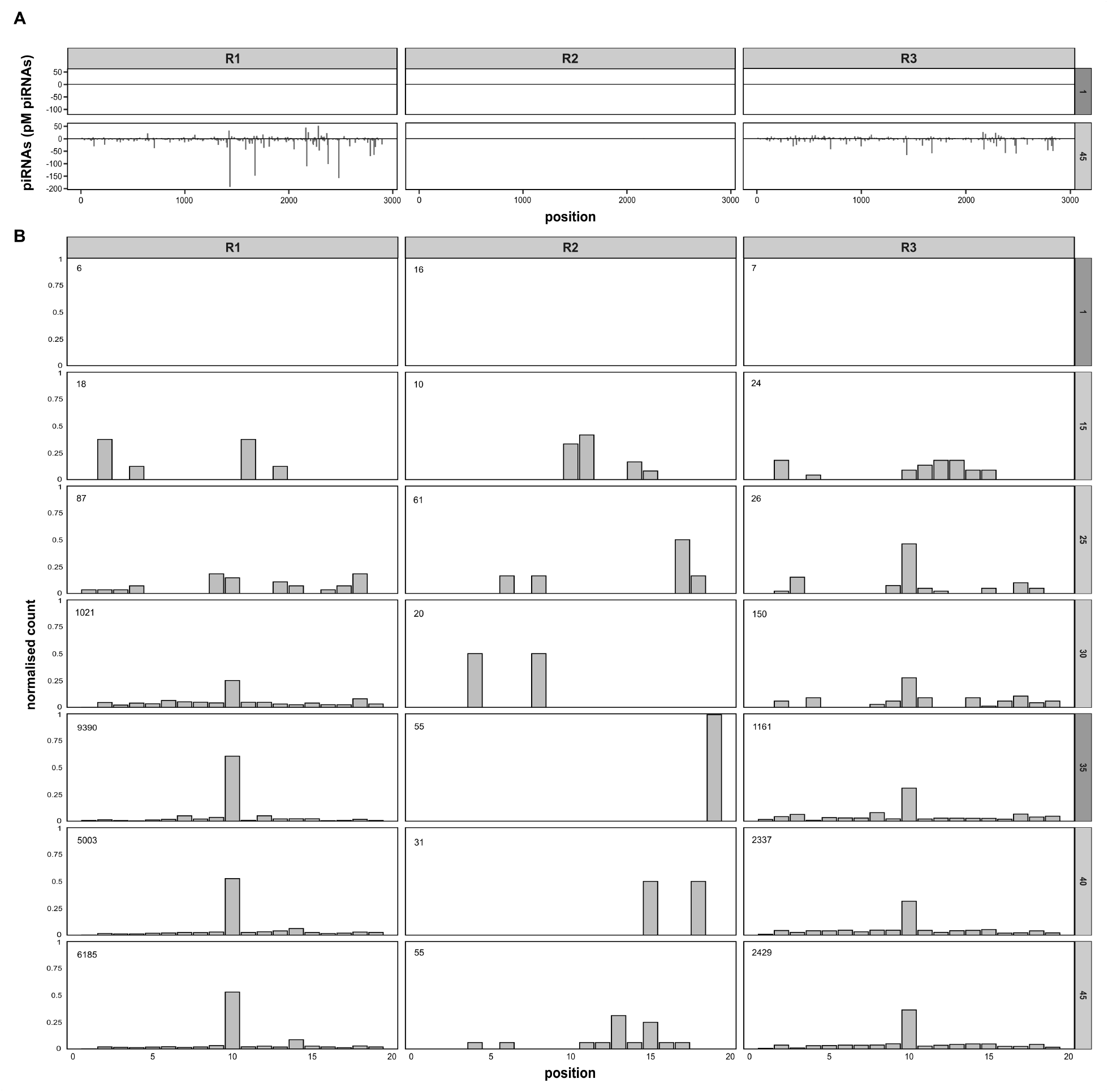
The low abundance of *P-element* piRNAs in R2 may be due to an inactive ping-pong cycle. **A** Distribution of piRNAs on the *P-element* at generation 1 and 45 (right panel). Sense piRNAs are shown on the positive y-axis and antisense on the negative. Samples from the whole-body of female flies are labelled in a light-grey and those taken from ovaries are in dark-grey. **B** Ping-pong signatures for the *P-element* in different replicates during the experiment (generations denoted in the right panel). Values in the top-left corner show the total number of *P-element* piRNAs in the sample.

### Rise of non-autonomous *P-element* insertions in R2

*P-element* activity can be regulated by non-autonomous insertions with IDs, such as the *KP*-element [7]. Proteins produced from such defective insertions may, for example, occupy available transposase binding sites, preventing mobilisation of the *P-element* [7, 56, 40]. *We posited whether the emergence of non-autonomous P-element* insertions, similar to the *KP*-element, could be responsible for the plateauing of *P-element* copy numbers in the absence of a piRNA-based host defence. To test this, we investigated the coverage and abundance of *P-element* IDs in each replicate over time (Fig. 3A). IDs of the *P-element* were detected using DeviaTE, which is based on split-reads (Fig. 3A). In both R1 and R3, *P-element* coverage broadly increased throughout the experiment. Although several IDs emerged (black arcs), a contiguous coverage across the entire *P-element* can be observed in R1 and R3, suggesting that abundant full-length insertions are present in these replicates (Fig. 3A). In contrast, coverage in R2 increased far more slowly (Fig. 3A). Several IDs in central regions emerged in R2 at early generations. By generation 63, central regions (positions 1000-1500), are almost completely devoid of coverage, suggesting that extremely few full-length insertions are present in R2 by this time (Fig. 3A). To substantiate this, we sequenced 11-12 individual flies from each replicate at generation 98 and estimated *P-element* copy numbers and IDs as described (Fig. 3B). Within replicates, *P-element* copies are fairly homogeneous (Fig. 3B). Furthermore, the individual flies from R2 have significantly lower copy numbers than those from R1 and R3, consistent with our pooled estimates (Fig. 1C). To determine whether full-length insertions are entirely absent in individuals, we next investigated the coverage at an individual level. We reasoned that if a region in the *P-element* has a coverage of zero, then the sample cannot contain a single full-length insertion (Fig. 3C, yellow regions). This approach is likely conservative, as non-overlapping IDs in different insertions could result in a contiguous coverage, despite the absence of full-length insertions in the sample. Analysis of individual coverage revealed that at least 6 of the 11 sequenced individuals from R2 lack full-length insertions, whereas we did not find a single sample without full-length in both R1 and R3 (Fig. 3C; Supplementary fig S8). Next, we investigated whether any *P-element* insertions with IDs could yield repressors of *P-element* activity, such as the *KP*-element [7]. Such repressors are characterised by two properties: i) transposase translation must be interrupted (due to deletions or premature stop codons) and ii) the DNA-binding domain, in ORF0, must be present (Fig. 4A,D) [42, 40, 34]. We aimed to assess the abundance of such putative repressors of *P-element* activity in our experimental populations. We used DeviaTE to identify the breakpoints of the IDs in each population (based on split-reads; Fig. 4B) and estimated the frequency of insertions with IDs (based on the number of split reads supporting an ID and the mean coverage). IDs with a frequency *<*0.05 were filtered out (Fig. 4C). We found internally deleted copies that may act as *P-element* repressors in both R2 and R3 (Fig. 4C). In R3, we found a single putative repressor at a low frequency (0.05), whereas two with higher frequencies were present in R2 (0.1 + 0.27). Additionally, it is not clear whether another abundant ID (0.24) in R2, in which ORF0 is truncated (potentially reducing DNA binding efficacy), also acts as *P-element* repressor. Hence, putative repressors account for an estimated 0% of the *P-element* insertions in R1, 5% of the insertions in R3 and between 37-61% of the insertions in R2 (Fig. 4C). By contrast, full-length insertions, encoding functional transposases, account for 58% of the insertions in R1, 50% of insertions in R3 and only 2% of the insertions in R2. Compared to the other replicates, R2 is characterised by just a few full-length insertions but a high number of copies with IDs that may act as repressors of *P-element* activity. We conclude that the plateauing of the *P-element* copy numbers in the absence of a piRNA-based host defence in R2 is likely due to the rapid emergence and proliferation of non-autonomous *P-element* insertions with properties similar to known repressors of *P-element* activity.

**Figure 3:**
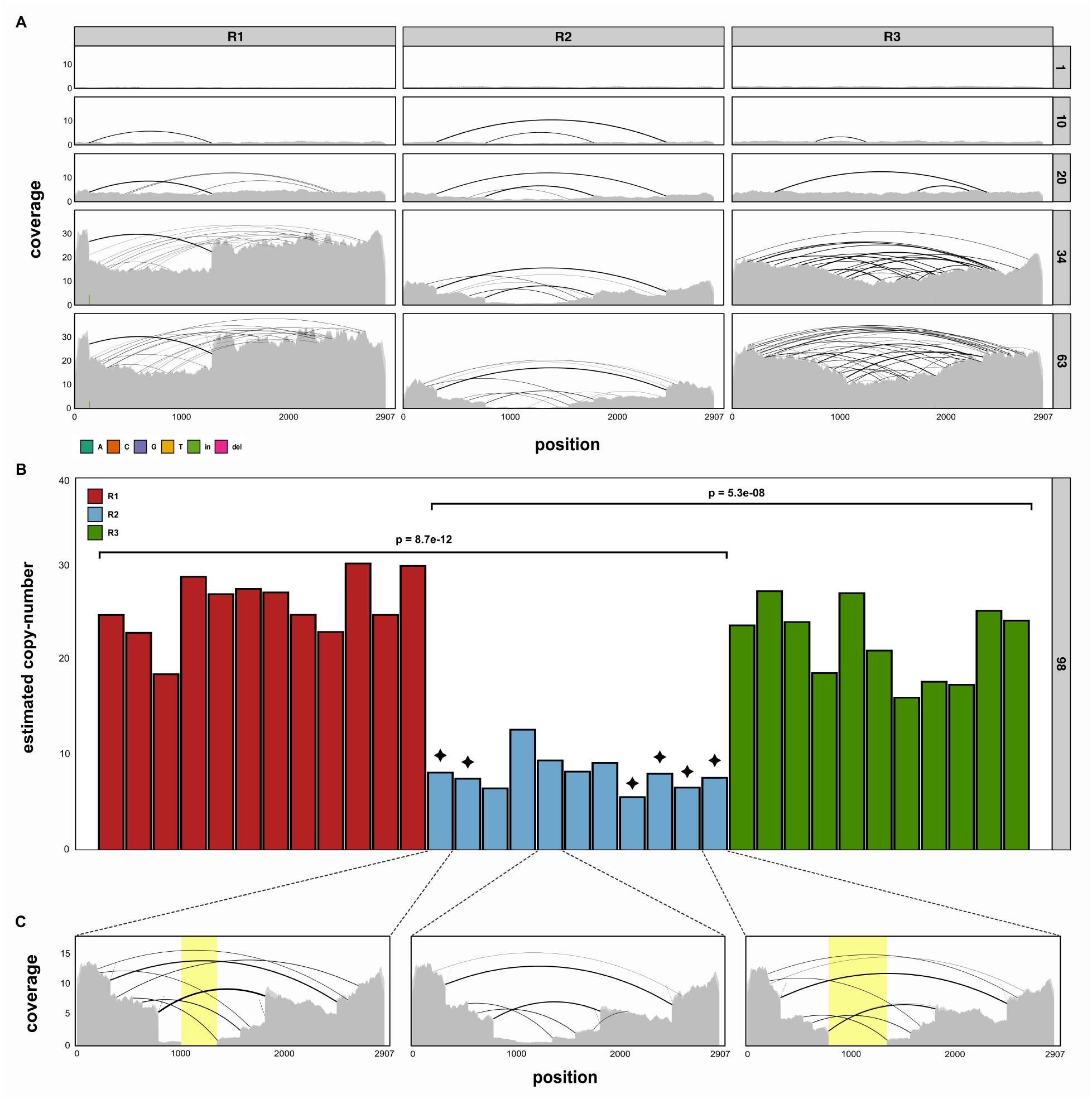
Overview of structural variation in the *P-element* during the experiment. **A** Abundance of *P-element* insertions and IDs across all three replicates (top) at different generations (right panel). Plots show the coverage normalised to single-copy genes, and the positions of IDs (inferred from split-reads) as arcs. **B** *P-element* copy numbers for 11-12 individual flies, sampled at generation 98. Significant differences between replicates are shown at the top (t-tests). Black stars denote individuals without a single full-length insertion. **C** *P-element* coverage of selected individuals from B. Yellow highlighted regions show areas with zero coverage. Individuals with zero-coverage regions cannot contain a single full-length *P-element* insertion.

**Figure 4:**
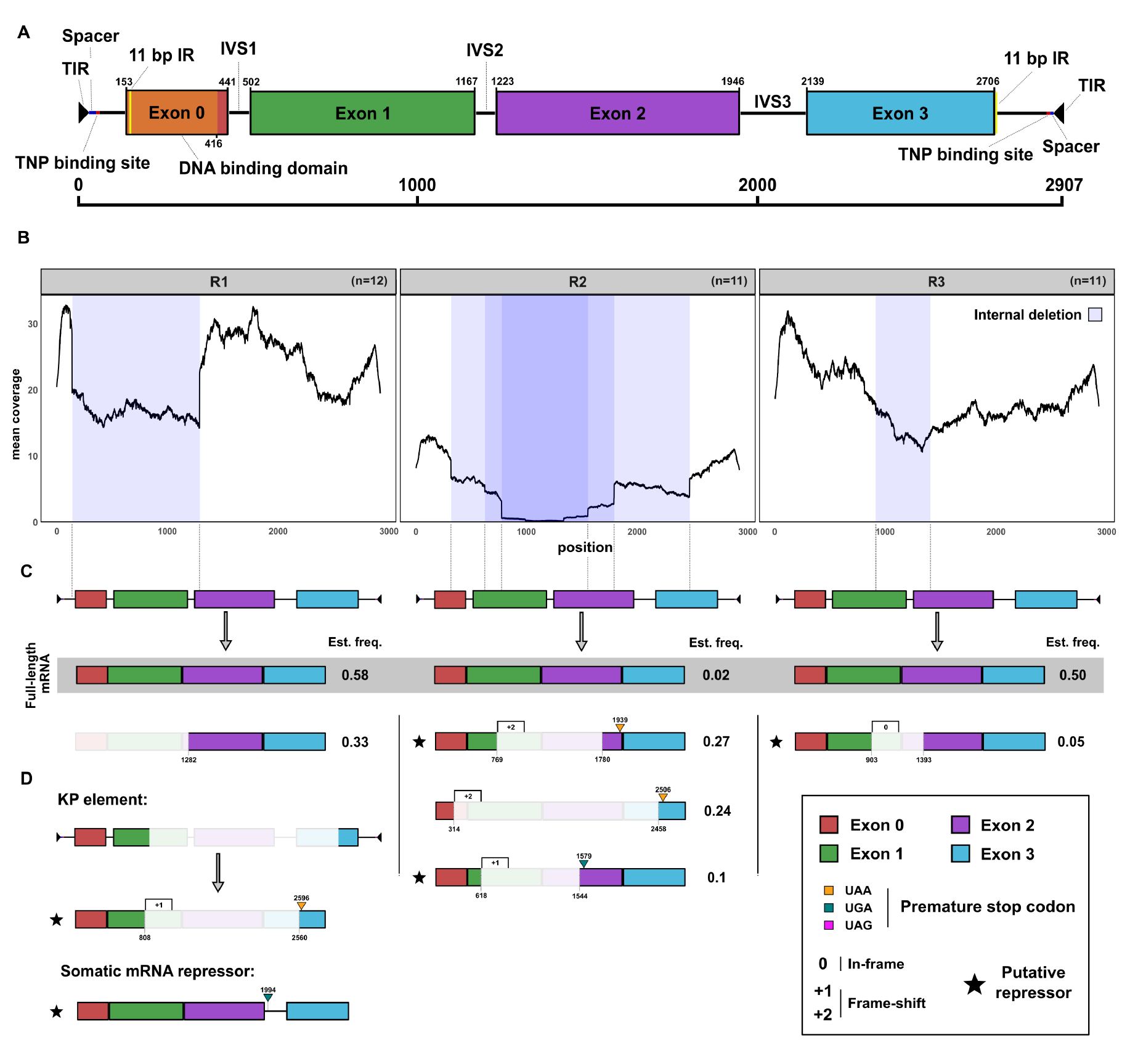
Proteins generated by *P-element* insertions with IDs in each replicate. **A** Schematic of the structure of the full-length *P-element*, highlighting the DNA binding domain and sites necessary for the mobilisation of the *P-element* (transposase binding sites, TIRs) [40, 42]. **B** Average coverage of the P-element across individuals in each replicate population, shaded blue regions denote prominent IDs (frequency ≥ 0.05).[73]. **C** Transposases encoded by different *P-element* insertions in our experimental populations. Their estimated frequency is shown on the right side. Premature stop-codons (yellow triangle) and frame shifts (rectangle with number) are highlighted. Putative repressors are highlighted by a star. **D** Schematic representation of the somatic mRNA repressor of the P-element and the *KP*-element (and its mRNA) [58, 7].

### Gonadal dysgenesis across experimental populations

The *P-element* was initially discovered as the cause of hybrid dysgenesis (HD) [29, 6], where crosses between males having the *P-element* with näive females displayed a wide range of different phenotypes, including atrophied ovaries, whereas reciprocal crosses (females with the *P-element* and näive males) do not. This non-reciprocity is due to piRNA-based host defences being maternally transmitted, while the *P-element* is transmitted by both parents [10]. Ovary atrophication is a result of germline stem cell arrest due to double strand breaks caused by *P-element* activity [47]. Atrophied ovaries thus provide an easily scored phenotypic indication of *P-element* activity (termed gonadal dysgenesis, GD).

We wanted to ascertain if GD, the hallmark of *P-element* activity, could be detected in our experimental populations. This enables us to test whether *P-element* activity has been reduced by mechanisms other than piRNAs in R2. We performed GD assays at 29°C, i.e. the temperature where *P-element* induced GD is most pronounced [29, 30]. For each cross, we used three sub-replicates of four males and four females, and then dissected ovaries of the F1. Crosses between the strong inducer strain, Harwich [29], and the *P-element* näive strain, DM68 (supplementary fig S1), acted as controls (Fig 5A). We expect strong GD for crosses between Harwich males and DM68 females, but no GD in the reciprocal crosses. From the intra-population GD assays, we detected little GD across all sub-replicates, indicating a low level of activity in all experimental populations (Fig 5B), further highlighting that *P-element* activity is low within the R2 population. Interestingly, levels of intra-population GD were consistently low throughout the experiment for R2, whereas GD-levels were initially low (until generation 15) for R1 and R3, but rose to ∼80% by generation 25, then dropped again to lower levels at generation 40 (supplementary fig. S9).

**Figure 5:**
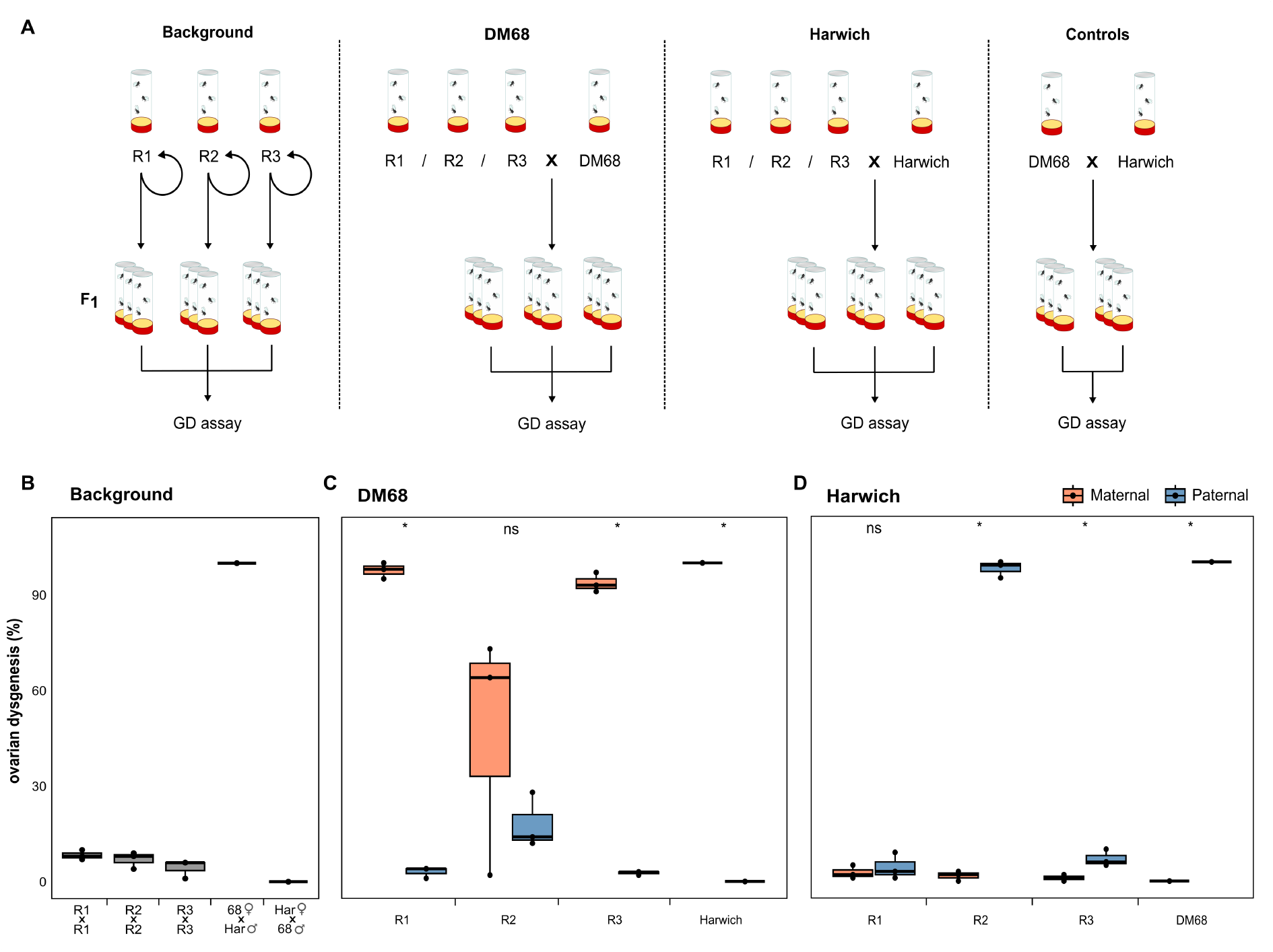
Gonadal dysgenesis (GD) assays suggest that the *P-element* is still weakly active in R2. **A** Schematic overview of crosses. All crosses were performed with flies from the experimental populations at generation 101. **B** Extent of intra-population GD in the experimental populations. As a control, reciprocal crosses among Harwich (with the *P-element*) and DM68 (without the *P-element*) are shown. **C** Crosses of males from the experimental populations to näive females (DM68) leads to high levels of GD (orange), while reciprocal crosses show minimal GD (blue). **D** Crosses of females of all experimental populations to Harwich males (a strong GD inducer strain with many *P-element* insertions [29]) induces GD in R2 but not in R1 and R3. Asterisks indicate significant differences in GD levels between reciprocal crosses (*p <* 0.05; paired t-test).

Next, we tested if *P-element* insertions in the experimental populations are able to induce GD, informing us as to whether they are still functional. We crossed males from the experimental replicates to the naïve females of DM68. As DM68 contains no *P-element* insertions and therefore no complementary piRNAs, their offspring will exhibit GD if males have sufficient numbers of functional *P-element* insertions. We found that crosses of males from all replicates with DM68 females induced GD, while reciprocal crosses did not (Fig. 5C). Interestingly, we see that the *P-element* in R2 could still induce a substantial amount of GD, albeit at a lower and more variable level than seen in R1 and R3. However, in R2 the reciprocal cross (R2 females with DM68 males) also has a slightly elevated GD level.

Lastly, we tested if the different replicate populations are able to silence the *P-element*, by crossing experimental females to Harwich. If females from the experimental populations have *P-element* piRNAs, we should expect little to no GD. Crosses with females from R1 and R3 did not exhibit GD consistent with the emergence of a piRNA-based host defence (Fig. 5D, 1D, 2). In contrast, crosses with females from R2 show strong GD (Fig 5D), indicating that a piRNA-based host defence against the *P-element* is still absent in R2 after over 100 generations of the experiment. Taken together, our GD assays suggest that an effective host defence (likely piRNAs) emerged in both R1 and R3, but not in R2. Nevertheless, R2 still contains functional *P-element* copies, which are able to induce GD. The absence of intra-population GD in R2 further suggests that the *P-element* activity is low in this replicate.

### Internally deleted *P-elements* seen in global populations

We sought to assess whether an absence of full-length insertions in individuals, as seen in R2 (Fig. 3), could also be observed in natural *D. melanogaster* populations. We investigated the frequency of different *P-element* insertions in publicly available data. Initially, we utilised a total of 753 short-read samples (strains or pooled populations) collected from all major continents [21, 64, 35, 57, 27, 53, 13]. The average normalised *P-element* coverage across different continents shows that coverage frequently decreases within central regions of the *P-element* (Fig 6A). This coverage dip is likely due to highly abundant IDs, such as the *KP*-element [7]. Based on this, we estimate that the fraction of samples containing at least one full-length insertion varies dramatically across continents. Our data suggests that full-length insertions of the *P-element* are rare in both Europe and Asia but more abundant in the Americas, Africa, and Oceania (Fig. 6A). However, this data needs to be treated with some caution, as we only considered the coverage in central regions of the *P-element* and as we have included pooled populations in our analysis.

**Figure 6:**
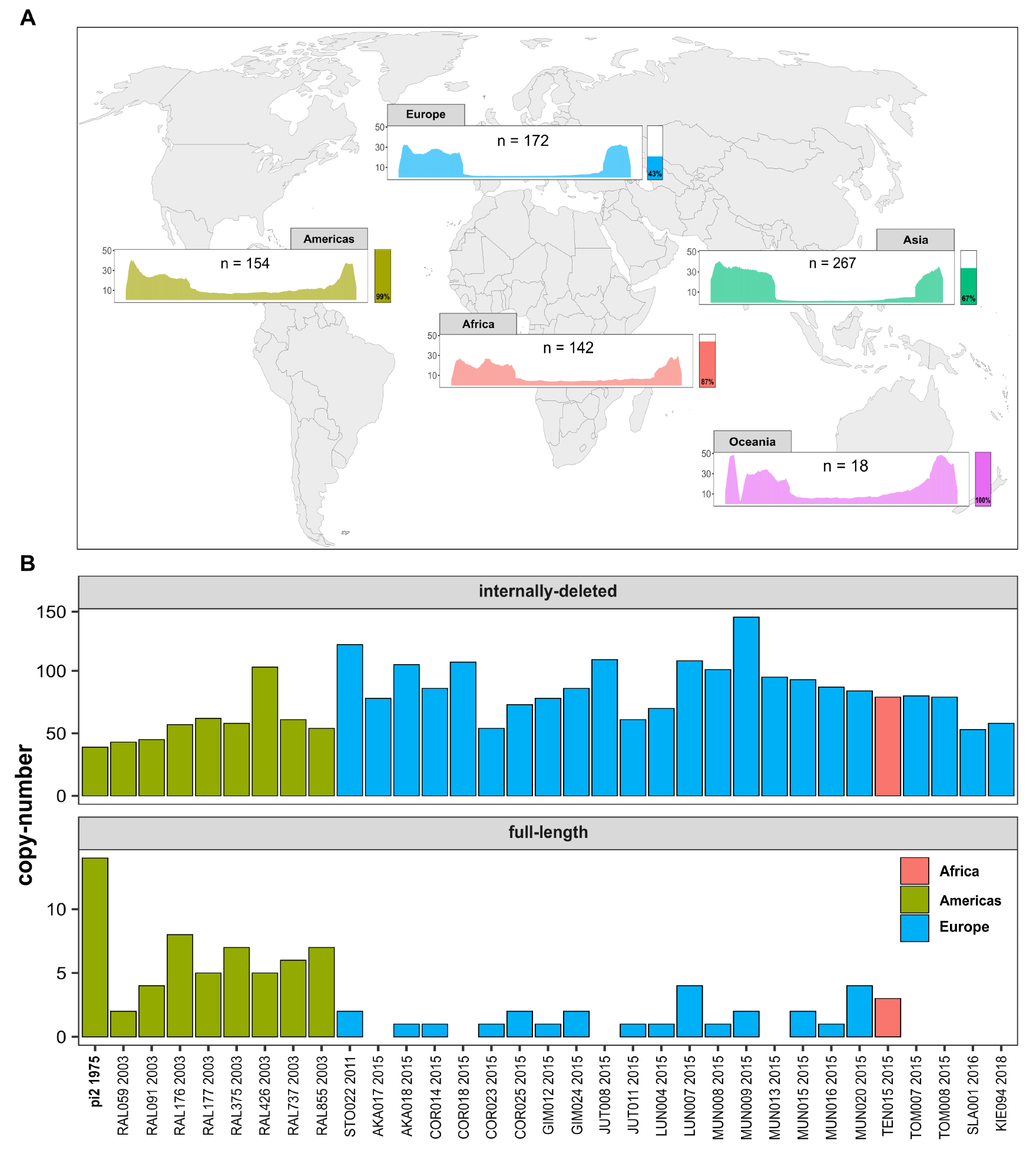
Abundance and structure of *P-element* insertions across natural *D. melanogaster* populations. **A** Mean coverage of the *P-element* in different geographic regions (data from 753 short-read datasets). Bars on the right provide a rough estimate of the fraction of samples containing at least a single full-length insertion. **B** Copy number of full-length *P-element* insertions and of insertions with IDs in long-read assemblies of recently collected *D. melanogaster* strains [57]. Strains are coloured by region and labels show strain name and collection year. pi2 (bold) is a frequently used inducer strain of GD, collected in 1975.

To further investigate the composition of the *P-element* in natural populations, we analysed 33 long-read assemblies of *D. melanogaster* strains, recently collected from Europe and North America [12, 76, 24, 57]. We used RepeatMasker to identify full-length insertions and insertions with IDs in these assemblies. All investigated strains contained *P-element* insertions (either full-length or insertions with IDs; Fig. 6B). In agreement with the short-read data, we found that full-length insertions were rare in strains from Europe but more abundant in strains from North America (Fig. 6A). Several of the European strains did not contain even a single full-length *P-element* insertion (Fig. 6B). This suggests that the absence of full-length insertions in populations invaded by the *P-element*, as observed in our R2, may also occur in natural *D. melanogaster* populations.

## Discussion

We introduced the *P-element* into three replicate populations of *D. melanogaster* and monitored the following invasion at the level of the genome and transcriptome for over 100 generations. We observed that copy numbers of the *P-element* stabilised at around 20-25 copies in two replicates (R1, R3), but at only ∼7 copies, in R2. Interestingly, copy numbers stabilised in R2 despite the absence of a piRNA-based host defence (until at least generation 45, from small RNA data). GD assays indicate that a piRNA-based host defence was still absent in R2 at generation 98 (females crossed with Harwich males induced GD; Fig. 5D). We found that non-autonomous *P-element* insertions with IDs rapidly emerged and proliferated in R2. Many individuals from R2 in later generations contain *P-element* insertions with IDs but are without a single full-length insertion. Several of these IDs share features of repressors of *P-element* activity, similar to the *KP*-element. We posit that the early appearance and propagation of non-autonomous *P-element* insertions is responsible for the stabilisation of *P-element* copy numbers in R2, despite the absence of a piRNA-based host defence. It has been suggested in previous works that non-autonomous elements may outcompete full-length insertions [54, 59, 34]. However, our work shows for the first time that non-autonomous insertions may emerge *de novo* within but a few generations in experimental populations and then proliferate to such an extent that TE copy numbers stabilise despite the absence of the host defence. Our work highlights a novel risk for TEs, i.e. that TE invasions can fail due to the emergence of non-autonomous elements.

This finding raises the question as to why non-autonomous *P-element* insertions spread in R2 in such a way that full-length insertions became so rare. An accumulation of *P-element* insertions with IDs could either be a consequence of positive selection or preferential mobilisation. Previous work has suggested that *P-element* insertions with IDs are likely preferentially mobilised [34, 33, 65, 26]. It is feasible that the shorter length of non-autonomous insertions facilitates a more effortless transposition. The proliferation of very short *P-element* insertions (Har-P) could also be responsible for the high rate of GD induced by some strains, such as Harwich [70]. A mobilisation advantage of non-autonomous elements has also been noted for other TE families [25]; in a direct competition, non-autonomous *Mariner* insertions were able to outcompete their autonomous counterparts [59].

Our work also leads us to consider why the piRNA-based host defence never established itself in R2. It is not yet clear what triggers the emergence of a piRNA-based host defence, but insertions in piRNA clusters, or siRNAs mediating the conversion of TE insertions into piRNA producing loci, have been suggested as potential mechanisms [41, 9, 4]. It is possible that *P-element* copy numbers in R2 were insufficiently abundant to trigger these mechanisms. In a previous study, we described another replicate population of *D. erecta*, wherein the host defence against an invading *P-element* also failed to be established [65]. Copy numbers of the *P-element* in *D. erecta* were over an order of magnitude higher than in this work (*D*.*ere* = 151, *D*.*mel* = 7) and acted to the severe detriment of the population’s fitness. This suggests that the mechanism triggering the host defence in *Drosophila* may not depend on TE copy numbers, as in plants [44]. In *D. erecta*, we also found multiple *P-element* insertions in piRNA clusters and a significant number of *P-element* siRNAs, suggesting that these two mechanisms are insufficient to trigger the establishment of the host defence [65]. Interestingly, non-autonomous *P-element* insertions with properties similar to the *KP*-element also proliferated in the unprotected *D. erecta* population [65]. This indicates that abundant *P-element* insertions with IDs might interfere with the establishment of a piRNA-based host defence. The mechanism by which this could be achieved remains unclear.

Another open question is why R2 males induced GD when crossed with naïve females but not with R2 females, despite both lacking a piRNA-based host defence (Fig. 5). We do not have a definitive answer to this, but we speculate that it could be linked to the dosage of non-autonomous elements. In intra-population crosses (e.g. R2), this is twice as a high as in crosses with naïve strains, and this higher dosage may be necessary to prevent *P-element* mobilisation. It is perhaps worth noting that the flies from R2 closely resemble the rare P’ strains [28]. These strains are able to induce GD (when crossed paternally to a naïve strain), yet at the same time are susceptible to GD (when crossed maternally to an inducer strain) [28]. This implies that P’ strains have active *P-element* insertions but no maternally transmitted piRNAs. One explanation is that the P’ phenotype is a transient stage, that can only observed for a few generations in strains actively being invaded by the *P-element*. Our findings raise the possibility that for some strains the P’ phenotype may be stable, wherein *P-element* activity might be controlled by non-autonomous insertions, *in lieu* of a piRNA-based host defence.

This study challenges our understanding of the evolutionary impact of failed invasions, where non-autonomous elements have proliferated at the cost of full-length insertions. Such failed invasions could pose a severe threat to the long-term sustained survival of TEs. TEs silenced by a host are likely to accumulate mutations over time, eventually resulting in the loss of functional copies within the host population [8, 63]. To persist, TEs must invade novel species, e.g. following horizontal transfer (HT). HT is likely a rare event, and TEs that fail to take advantage of the limited opportunities to spread into novel species may be unable to persist. The rapid proliferation of non-autonomous insertions, as observed in R2, poses a two-fold threat to the prolonged existence of TEs. First, abundant IDs could effectively ‘immunise’ a species (or population) to further invasions from a TE. Any newly introduced full-length insertions (e.g. recurrent HT) may be quickly outcompeted by the non-autonomous insertions already pervasive in the species. Second, species with abundant non-autonomous elements are likely not ‘infective’. Due to the scarcity of full-length insertions, HT from populations with abundant non-autonomous insertions to naïve species is unlikely to trigger a TE invasion. Populations with abundant non-autonomous insertions are likely evolutionary dead-ends: resistant to further invasions, yet unable to infect other species. Consistent with previous works, we show that a proliferation of non-autonomous *P-element* insertions can also be observed in some natural populations of *D. melanogaster*, where, especially those from Europe, few have full-length insertions (Fig 6 [74, 7]). These European populations could exhibit the two-fold cost of the proliferation of non-autonomous elements, threatening the long-term persistence of TEs.

## Materials and Methods

### Experimental populations

We introduced the *P-element* into DM68, a *D. melanogaster* strain collected 1954 in Israel, via micro-injection of the plasmid ppi25.1 (kindly provided by Dr. Erin Kelleher). Injections were performed by Rainbow Transgenic Flies Inc (https://www.rainbowgene.com/; Camarillo, CA, USA). We obtained 7 lines containing the *P-element* by crossing transformed adults (2 males and 3 females). Transformed lines were maintained at 20°C for 3 generations before setting up the experimental populations.

To establish the experimental populations, we crossed five males from five *P-element* containing lines with 5 naïve virgin females and allowed them to mate for 3 days. After mating, we mixed these 50 flies [(5M+5F)*5] from the crosses with 200 naïve *D. melanogaster* flies. We maintained 3 replicates of the experimental populations with a population size of *N* = 250 for over 100 generations at 25°C using non-overlapping generations.

### Genomic analysis

For genomic sequencing, we sequenced pools of 60 flies using Illumina 2×125bp reads. The individual flies at generation 98 were sequenced by BGI with 150bp reads (BGI Tech Solutions, Hong Kong). The abundance of the *P-element* was estimated with DeviaTE. Illumina short reads were aligned to a list of the consensus TE sequences in *D. melanogaster* (https://github.com/bergmanlab/drosophila-transposons [55]), alongside three single-copy genes; *tj, RpL32* and *rhi* (FlyBase release 2017 05). Coverage of the TEs was normalised to the abundance of the single-copy genes to estimate the abundance of the TE. DeviaTE was also used to obtain information about SNPs and indels within the *P-element*.

### Transcriptomic analysis

RNA was collected and sequenced from 30 female flies, either from whole-fly tissue or ovaries. Small RNA and RNA from these samples was sequenced by Fasteris (https://www.fasteris.com/en-us/) and BGI (BGI Tech Solutions, Hong Kong). RNA samples were treated with DNase and poly-A selected before they were sequenced using Illumina 2×100bp reads (NovoSeq). RNA data were aligned using GSNAP (version 2014-10-22; [77]) to the reference of *D*.

*melanogaster* (r6.52; Flybase) combined with the consensus sequences of TEs in *D. melanogaster* ([3]). The coverage and the splicing level of the *P-element* were visualized in R. Adaptor sequences of the small RNA data were removed with cutadapt (v2.6 (Martin, 2011)). We aligned the small RNA data to the *D. melanogaster* transcriptome (r6.62, Flybase) combined with the consensus sequences of TEs using novoalign (v3.09.00; http://www.novocraft.com/). The abundance of piRNAs, the distribution of piRNAs within the *P-element*, the length distribution of the piRNAs, the ping-pong and phasing signature were computed using previously described Python scripts [34, 65].

### Properties of *P-element* insertions with IDs

We investigated the most abundant IDs in all replicate populations using the combined data from individual flies sequenced at generation 98. ID positions were inferred from split-reads, aligned by DeviaTE (see above). Frameshifts and premature stop codons were identified using ORFfinder [60] and Expasy [16]. We estimated the frequency of the IDs based on the count of split-reads relative to the coverage. First for each replicate we calculated the average coverage of *P-element* regions not covered by IDs, excluding 50 bp at either end to avoid lower coverage regions. Next, we computed the proportion of each individual ID in the populations as the count of split-reads divided by the mean coverage outside of regions with IDs. Frequencies of the full-length germline mRNA were estimated as the minimum coverage of the *P-element* across all individuals of a replicate, again divided by the mean coverage outside of regions with IDs.

### Gonadal dysgenesis

Gonadal dysgenesis assays were set-up using 3 sub-replicates, with the exception of the intra-population assays conducted during the experiment where a single replicate was used (supplementary fig. S9). To estimate the level of GD for each cross we allowed 4 virgin females and 4 males to mate for 2 days. Selected flies were placed in cages and left to lay eggs for 2 days. The remaining eggs were kept at a constant 29^°^C until flies eclosed. The females were then taken for dissection. For each sub-replicate, we dissected 50 flies (100 ovaries) in 1x PBS solution and scored the proportion of atrophied ovaries.

### *P-element* in natural populations

We gathered a total of 753 publicly available short-read datasets ([21, 64, 35, 57, 27, 53, 13]) and 33 long-read assemblies ([12, 76, 24, 57]). For the short-read data, we estimated the abundance of the *P-element* with DeviaTE, as described above. We assumed that samples with a normalised coverage *>* 1 over the whole sequence have at least one full-length insertion. To analyse *P-element* composition in the long-read assemblies, we used RepeatMasker [69] (open-4.0.7; -no-is -s -nolow) with a custom library that included only the *P-element* consensus sequence. Samples with at least one insertion with a length *>* 2325 bp (80% of the *P-element* with 2907 bp) are considered to contain a full-length insertion.

## Data access

The sequencing data generated in this work are available from NCBI (BioProject ID: PRJNA1198884).

All analysis performed here is available at: https://github.com/divygenome/Dmel_Pelement_Invasion

## Author contributions

R.K. conceived the project. D.S. and M.B. performed the experiments. M.B., D.S. and R.P. analysed the data. R.K and M.B. wrote the manuscript.

## Acknowledgments

We thank Erin Kelleher for providing ppi25.1, Almorò Scarpa and all other members of the Institute of Population Genetics for their feedback and support.

